# Molecular Magnetic Resonance Imaging of Dysregulated Zinc Secretion Detects Early Pancreatic Ductal Adenocarcinoma Lesions and Response to KRASG12D Inhibitor Treatment

**DOI:** 10.1101/2024.09.15.613145

**Authors:** Veronica Clavijo Jordan, Mozhdeh Sojoodi, Farzad Moloudi, Patricia Gonzalez-Pagan, Manyu Jin, Pamela Pantazopoulos, Ethan French, Jonah Weigand Whittier, Nicholas Rotile, Mehrad Tavallai, Jill Hallin, Ian Ramsay, Eric M Gale, Stephen C. Barrett, Nabeel El-Bardeesy, Motaz Qadan, Kenneth K. Tanabe, Peter Caravan

**Affiliations:** Martinos Center for Biomedical Imaging, Department of Radiology, Massachusetts General Hospital, Harvard Medical School, Charlestown, MA, USA; Institute for Innovation in Imaging (i_3_), Massachusetts General Hospital, Harvard Medical School, Boston, MA, USA; Division of Gastrointestinal and Oncologic Surgery, Massachusetts General Hospital, Harvard Medical School, Boston, USA; Mirati Therapeutics, Inc., San Diego, California, USA; Massachusetts General Hospital Cancer Center, Harvard Medical School, Boston, MA, USA; The Cancer Program, Broad Institute, Cambridge, MA, USA

## Abstract

Pancreatic ductal adenocarcinoma (PDAC) is a highly lethal cancer, primarily due to late-stage diagnosis and limited treatment options. Zinc homeostasis is markedly dysregulated in PDAC and this dysregulation can be probed by administering a secretagogue to stimulate zinc secretion (SSZS) in the exocrine pancreas and imaging this secretion with a zinc sensitive MRI probe. This study demonstrates the potential of SSZS MRI for early detection, monitoring treatment response, and assessing recurrence after treatment withdrawal in PDAC. Our approach relies on interrogating the pancreas, circumventing the challenge of locating small, elusive tumors. By SSZS MRI, we detected PDAC by observing the unique zinc hypersecretory activity of the pancreas when malignancy is present. We observed dysregulation of zinc transporters in both human and mouse pancreas containing PDAC and confirmed secretagogue-stimulated zinc secretion in vitro and in vivo. We found that combining secretagogues such as secretin and caerulein maximized zinc secretion and as such MRI signal in the pancreas. Notably, SSZS MRI detected treatment responses to KRAS G12D inhibition within 3-5 days and identified cancer recurrence as early as one day post-treatment withdrawal. Additionally, secretagogue stimulation improved treatment responses and delayed recurrence in both treatment models. These findings suggest that SSZS MRI could significantly enhance PDAC diagnosis and management, providing a novel, non-invasive imaging modality to improve patient outcomes.

**STATEMENT OF SIGNIFICANCE:** This study demonstrates the utility of secretagogue-stimulated zinc secretion (SSZS) MRI in detecting pancreatic ductal adenocarcinoma (PDAC) at early stages, monitoring treatment responses, and assessing cancer recurrence, thereby offering a promising non-invasive imaging modality to improve PDAC patient management and outcomes.

## INTRODUCTION

Pancreatic ductal adenocarcinoma (PDAC) is the predominant form of pancreatic cancer, notorious for its dismal prognosis, with a 5-year survival rate of less than 12%.(1) The insidious onset and asymptomatic progression of PDAC mean that most patients present with unresectable, locally advanced, or metastatic disease at the time of diagnosis.(2) Surgical intervention, such as pancreaticoduodenectomy or distal pancreatectomy, remains the sole curative option, but unfortunately, fewer than 20% of patients qualify for surgery. (3,4) Strikingly, patients diagnosed with lesions smaller than 10 mm can achieve a 5-year survival rate as high as 60%, (5,6) highlighting the critical need for early detection. Despite successful resection, 80% of patients experience recurrence within five years.(4)

The current diagnostic approaches rely on serum biomarkers, imaging, and biopsy sampling. Biomarkers like cancer antigen 19-9 (Ca19.9) have low sensitivity and specificity. Non-molecular imaging techniques like CT and MRI offer moderate sensitivity but they either involve exposure to ionizing radiation or lack the molecular information to render them specific to cancer.(6,7) Although magnetic resonance cholangiopancreatography (MRCP) provides excellent ductal visualization, its diagnostic utility is limited without quantitative molecular information to differentiate clinically significant lesions.(8) Nevertheless, imaging techniques are used to locate the tumor and perform invasive biopsy sampling to confirm disease by histology. Thus, an imaging technique integrating functional pancreatic biology with the non-invasive benefits of MRI could be invaluable.

Current therapeutic approaches for those not eligible for surgery typically involve chemoradiation aimed at downstaging tumors for resection or controlling the disease. Regimens such as gemcitabine-based therapies and FOLFIRINOX (a combination of folinic acid, fluorouracil, irinotecan, and oxaliplatin) are standard but come with significant cytotoxicity and side effects, making early determination of treatment efficacy crucial. Furthermore, despite recent advancements, such as KRASG12D mutation inhibitors targeting the KRAS oncogene prevalent in ∼40% of PDAC patients,(9) effective alternative therapies remain limited.

The pancreas performs both endocrine and exocrine functions. In exocrine tissue, acinar and ductal cells work together to secrete digestive enzymes and bicarbonate necessary for digestion. (10) It has been hypothesized that PDAC arises from transdifferentiation of acinar cells into ductal structures in response to specific insults, such as tissue damage, inflammation, or mutation in genes such as KRAS, p16/CDKN2A, TP53, and DPC4/SMAD4. (11,12) A significant hallmark of glandular cancers, including PDAC, is the dysregulation of zinc homeostasis.(13) Zinc plays a crucial role as co-factor for various enzymes and transcription factors and acts as a secondary messenger in several signaling pathways. Proper intracellular and extracellular zinc concentrations are tightly regulated, and any dysregulation can contribute to the initiation and progression of PDAC. (4)

Zinc homeostasis is maintained primarily by the regulated actions of zinc transporters located on the cell membrane. (14) Mammalian zinc transporters are classified into two major families: the ZIP family (SLC39A proteins), which facilitates zinc influx into the cytoplasm from the extracellular environment, and the ZnT family (SLC30A proteins), which exports intracellular zinc. (15) Various studies have implicated the dysregulation of zinc homeostasis and transporter expression in the transformation from healthy to malignant and invasive PDAC. Notably, the upregulation of zinc import transporters ZIP3, and ZIP4 has been linked to the progression of pre-malignant pancreatic intra-epithelial neoplasia (PanIN) cells to invasive adenocarcinoma (16–18). Therefore, we hypothesize that measuring dysregulated zinc homeostasis in PDAC could serve as an indicator of PDAC presence and allow monitoring of treatment efficacy or recurrence.

It was shown that a gadolinium-based MR probe could detect changes in zinc flux noninvasively.(19,20) The Zn probe, GdL, comprises a stable Gd-DOTA like chelate and a pendant bis(pyridyl) group can bind to zinc with an affinity of K_D(Zn)_ = 2.35 μM. When the probe binds zinc, the resultant complex has high affinity for serum albumin. Binding to serum albumin has two beneficial effects: it serves to increase the relaxivity and in turn increase MR signal and it also localizes the probe in regions of high Zn concentration.(21) We used a bolus of glucose as an exogenous secretagogue to stimulate the secretion of zinc from intracellular stores and were able to detect zinc co-secretion with insulin as a result of increases in blood glucose in the endocrine pancreas using MRI with GdL, a technique termed secretagogue-stimulated zinc secretion MRI (SSZS-MRI). (20,22–24) SSZS-MRI was also demonstrated in the prostate, again using glucose as a secretogogue. The healthy prostate has the highest zinc levels in the body and these levels are markedly reduced in prostate cancer, (25) we previously demonstrated that the onset of prostate cancer was detectable as a decrease in the SSZS-MRI signal. (26,27) Here we propose to use SSZS-MRI in the exocrine pancreas.

Functional receptors that mediate zymogen granule secretion in the exocrine pancreas have been identified for secretagogues such as caerulein (cholecystokinin), secretin, acetylcholine, gastrin releasing peptide, substance P, and vasoactive intestinal peptide.(28) These secretagogues act on their respective G-protein-coupled receptors (GPCRs) located in the basolateral membrane and stimulate the secretion of zymogen granules docked on the luminal side. GPCRs act via Ca^2+^ movement triggering depolarization-stimulated exocytosis, or by activation of adenylate cyclase which in turn rises cellular cAMP activating exocytosis through cAMP-mediated phosphorylation of protein kinase A.(28–30) Several reports in animal models and in patients indicate that a combination of both pathways potentiate an even greater zymogenic secretion.(28–32) In the current study, we use the exocrine secretagogues, caerulein and secretin to evoke zinc exocrine secretion in the healthy and tumor-bearing pancreas. In an orthotopic syngeneic mouse model of PDAC we evaluate the effectiveness of SSZS-MRI in detecting zinc dysregulation at the onset and throughout the progression of pancreatic cancer. We also assess the capability of SSZS-MRI to monitor the response to KRASG12D treatment and to detect cancer recurrence after the treatment is stopped. Our primary objective is to introduce a novel, accurate method for detecting malignant PDAC lesions in the pancreas, aiming to transform early diagnostic and therapeutic monitoring practices in pancreatic cancer.

## RESULTS

### Dysregulated zinc homeostasis in human and mouse PDAC

Microscopically, human PDAC is formed by atypical tubular glands resembling medium-sized or smaller pancreatic ducts embedded in fibrotic stroma consisting of stromal cells, inflammatory cells, and extracellular matrix proteins. (11) Ductal obstruction caused by PDAC lesions results in inflammation in the adjacent tissue characterized by loss of acinar cells and small metaplastic ductal structures. (33) Six human PDAC specimens were collected, and three major regions were classified: PDAC lesions, pancreas tissue surrounding PDAC, and distal healthy pancreas (**Figure 1 A-C)**. Zinc import and export transporters ZIP3, ZIP4, and ZNT1 expression was evaluated in all three pancreas tissue regions. We found significant increase in ZNT1 expression in PDAC adjacent pancreas tissue versus distal normal pancreas (46 ± 14 % vs. 21 ± 6 %). Similarly, the expression of ZIP3 and ZIP4 was elevated in PDAC, and surrounding pancreas tissue compared to distal healthy pancreas (ZIP3, PDAC 35 ± 26 and surrounding pancreas 47± 35, vs. healthy pancreas 19 ± 16,) and (ZIP4, PDAC 40 ± 20 and surrounding pancreas 52± 25, vs. healthy pancreas 14 ± 9). Our data showed upregulation of zinc importers and exporters in PDAC and in surrounding PDAC pancreas tissue versus distal healthy pancreas.

**Figure 1.**
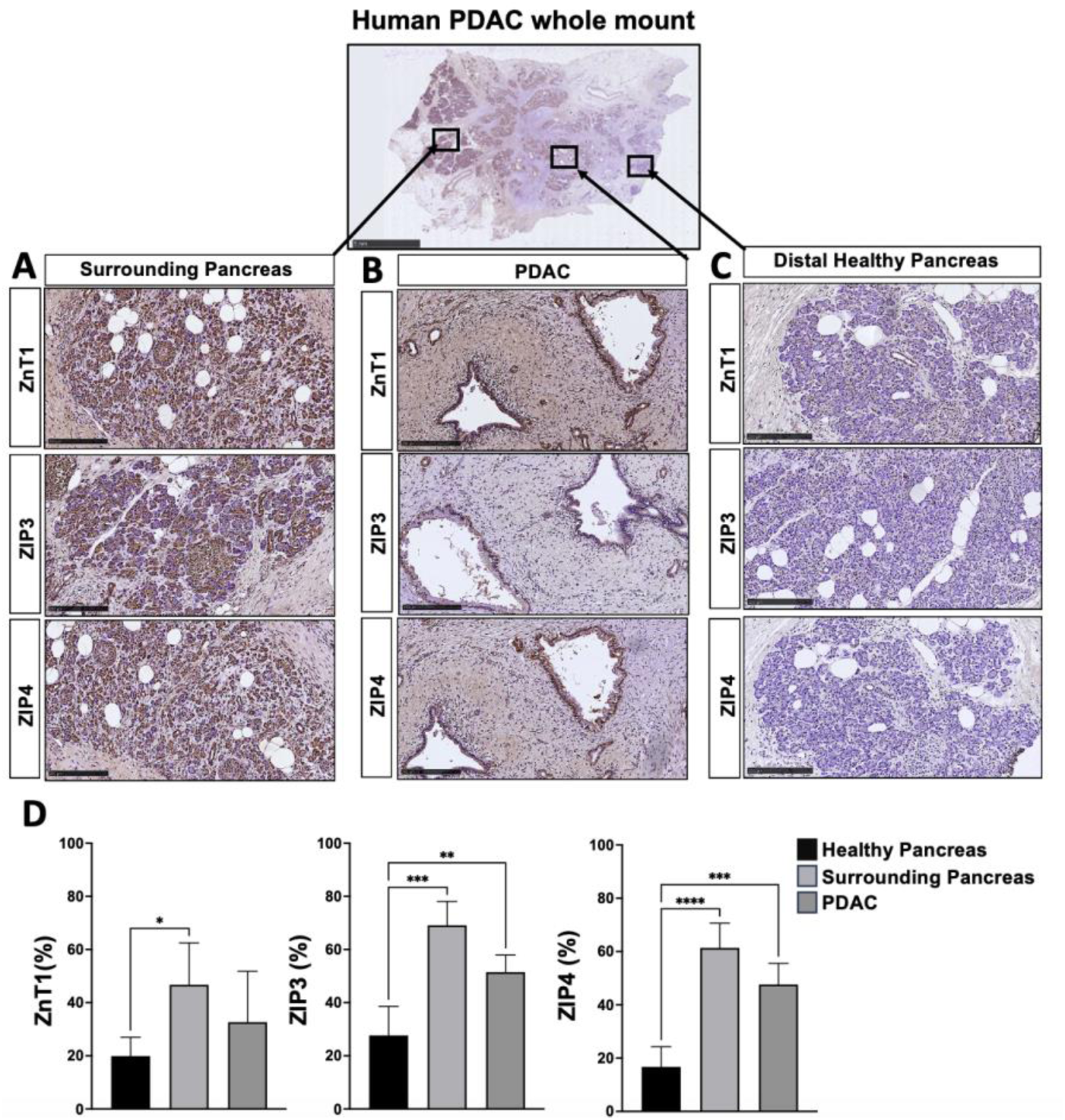
Dysregulation of zinc transporters in human PDAC. Immunohistochemistry staining showing expression of export transporter ZnT1, and import zinc transporters ZIP3, and ZIP4 for **A)** pancreas tissue surrounding PDAC, **B)** PDAC lesions, and **C)** distal healthy pancreas. (N = 6) Error bars are SD. Statistical significance was determined by one-way ANOVA with Tukey test for multiple comparisons. *p<0.05, ** p < 0.01, *** p <0.001, **** p<0.0001.

### The tumor bearing pancreas exhibits stimulated zinc secretion *in vivo*

To determine if we can monitor zinc release from intracellular stores *in vivo*, we used a previously reported cell-impermeable Zn sensistive MRI probe (20). The mechanism of zinc detection in vivo is illustrated in **Figure 2A** where the pancreas exhibits high contrast only after co-administration of the Zn probe and an exogenous exocrine secretagogue. When the secretagogue is administered the zinc probe binds to the secreted zinc and then forms a complex with albumin, this macromolecule exhibits magnetic properties that confer high contrast in the stimulated pancreas by T_1_-weighted MRI. To show this mechanism, PDAC mice were generated by orthotopic injection of syngeneic KPC cells (Hy15549) directly into the tail of the pancreas as previously described (34), and illustrated in **Figure 2B (left)**. In this model, mice develop progressive disease starting from low grade pancreatic intraepithelial neoplasia (PanINs), as early as 1-week post implantation, to multinodular tumors after 2-weeks (**Figure 2B, right**). To observe the effect of secretagogue stimulation in early and advanced stages of PDAC, we allowed PDAC cells to grow in the pancreas for 2-weeks while performing SSZS-MRI 1-week and 2-weeks after tumor cell implantation. We administered 10 μg/kg caerulein or saline i.p. as control to fasted healthy mice and PDAC mice followed by 0.07 mmol/kg Zn probe i.v. 5 minutes later. MR images of the healthy and tumor-bearing pancreas before and 6 minutes after caerulein are shown in **Figure 2C**. The pancreas of animals bearing tumors for 1-week after receiving caerulein show a significant increase in contrast relative to those receiving saline (**Figure 2C middle row**), and this signal increase is higher than that seen in the pancreas of healthy mice (**Figure 2C top row)**. By quantifying the change in contrast to noise ratio (ΔCNR) in the pancreas we see that the ΔCNR over the entire period (area under the curve) after caerulein stimulation relative to its saline control is largest for animals bearing a tumor for 1 week versus 2 weeks (caerulein 1w: 205 ± 29 ΔCNR *min vs. saline 1w: 31 ± 14 ΔCNR *min, p< 0.001 and caerulein 2w: 146 ± 29 ΔCNR *min vs. saline 2w: 117 ± 20 ΔCNR *min, p = 0.88). Similarly, we performed SSZS MRI in a second animal model where syngeneic Han4.13 PDAC cells are injected into the pancreas of FVB-strain mice (**Figure 1SA**). Consistent with results in animals of C57BL6 background, the FVB mice showed a relative increase in SSZS MR image enhancement in the pancreas of animals receiving caerulein versus saline. Similarly, animals bearing a tumor for 2 weeks showed no difference in SSZS MR image enhancement in the pancreas of animals receiving caerulein or saline (**Figure 1SB**).

**Figure 2.**
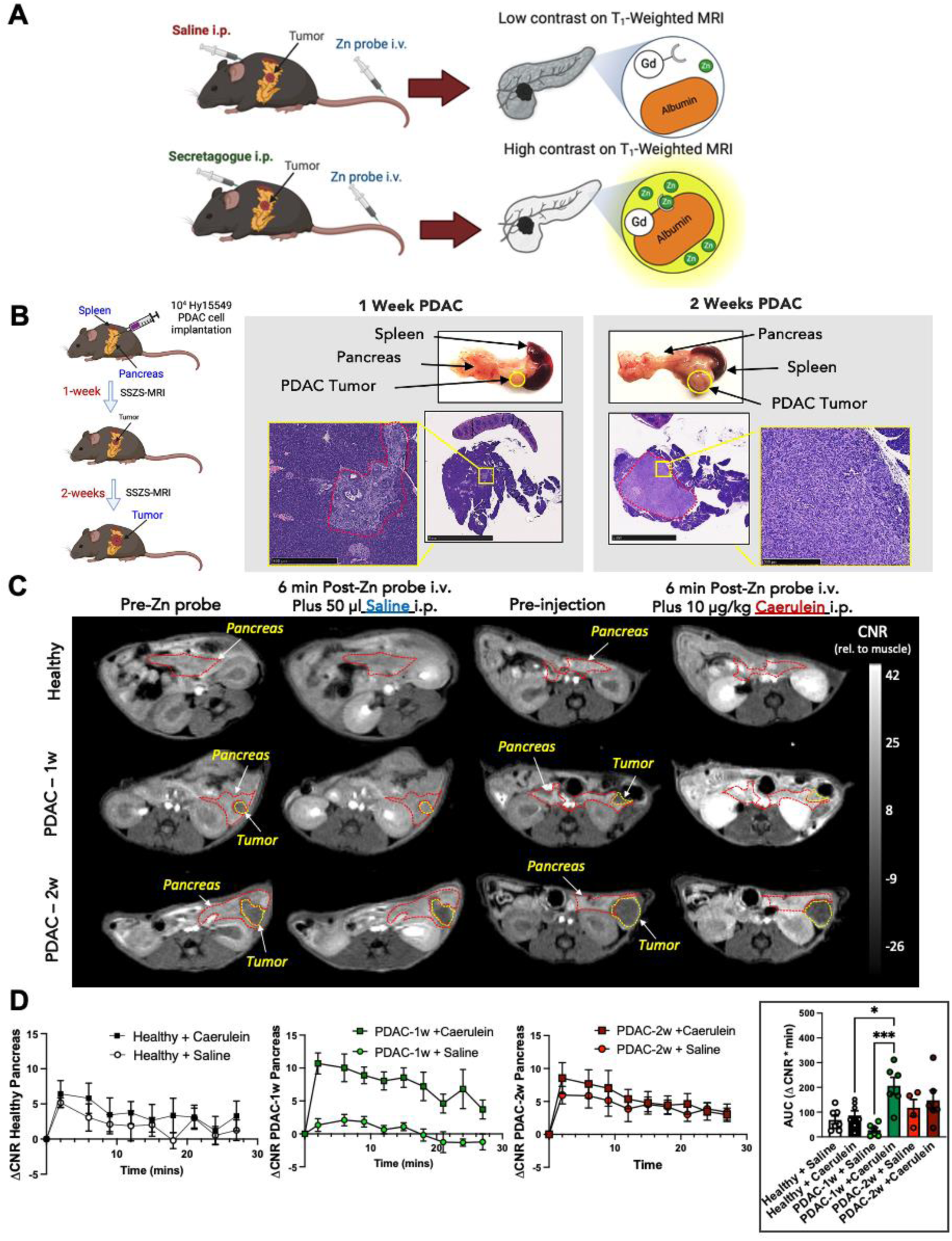
Increased stimulated zinc secretion in the tumor-bearing pancreas detected by MRI. **A)** Mechanism of contrast generated by secretagogue stimulated zinc secretion (SSZS) MRI in PDAC-bearing animals. **B)** (*left*)Surgical and imaging schedule to evaluate secretagogue stimulated zinc secretion MRI in early and advanced orthotopic PDAC model, SSZS -MRI evaluation is performed 1- and 2-weeks after cells are implanted into the pancreas. (*right*) Gross anatomy images and H&E staining illustrate the size, and location of the 1-and 2-week tumor relative to the spleen. **C)** *In vivo* MRI of a healthy mouse, 1-week, and 2-weeks post orthotopic PDAC tumor implantation. Representative T_1_-weighted MR Images were collected before and 6 minutes after 0.07 mmol/kg Zn probe i.v. injection. **D)** Change in contrast to noise ratio (△CNR) of a region of interest encompassing the pancreas relative to a ROI of muscle within the same imaging slice. The healthy pancreas does not exhibit stimulated zinc secretion compared to the PDAC-bearing pancreas. *Inset:* Area under the curve of △CNR vs. time for all groups. Error bars are SEM. Statistical significance was determined by one-way ANOVA with Tukey test for multiple comparisons. *p<0.05, ** p < 0.01, *** p <0.001, **** p<0.0001.

### Multi-secretagogue synergy maximizes zinc secretion in the pancreas

Secretin is a known exocrine secretagogue that stimulates secretion of bicarbonate in the pancreas by interacting with pancreatic ductal cells, (35) while caerulein stimulates the secretion of pancreatic enzymes by interacting with acinar cells.(36) Here we stimulated animals with each secretagogue individually and as a cocktail to maximize the secretion from both acinar and ductal cells in the pancreas. **Figure 3A** shows T_1_-weighted MR images of mice bearing tumors for 1 and 2-weeks before and 6 minutes after Zn probe and secretin or cocktail (secretin+caerulein) administration. Serial T_1_-weighted images were acquired over 27 minutes and ΔCNR vs time traces were calculated. **Figure 3B** shows the complete comparison of ΔCNR vs time and its AUC for Caerulein, Secretin, Cocktail, and Saline groups together. In the 1-week PDAC model, caerulein induced a markedly higher secretion compared to secretin in the tumor-bearing pancreas, signifying a robust acute secretory response that can be captured by MRI. This enhanced secretion is evident through a significantly greater AUC for caerulein and cocktail versus saline (caerulein: 205 ± 29 ΔCNR *min. vs. saline: 32 ± 14 ΔCNR *min. p<0.001 and cocktail: 185 ± 32 ΔCNR *min, vs. saline: 32 ± 14 ΔCNR *min. p<0.05), while secretin did not elicit a large secretory response compared to saline (secretin: 45 ± 24 ΔCNR *min vs. saline: 32 ± 14ΔCNR *min. p>0.05) indicating that caerulein may be more sensitive at detecting early PDAC lesions. Interestingly, in the 2-week PDAC model, secretin and the combination (caerulein + secretin) cocktail showed the highest secretory response (secretin: 287 ± 17 ΔCNR *min vs. saline: 117 ± 20 ΔCNR *min, p< 0.05 and cocktail: 317 ± 33 ΔCNR *min vs saline: 117 ± 20 ΔCNR *min, p< 0.05). These results support the hypothesis that the secretory capacity of the pancreatic tumor microenvironment changes over time and is particularly responsive to caerulein or cocktail stimulation at the nascent stage of tumor growth. This observation may reflect acinar-to-ductal metaplasia resulting in more ductal-like cell populations in more advanced PDAC affecting its secretory response to different stimuli.

**Figure 3.**
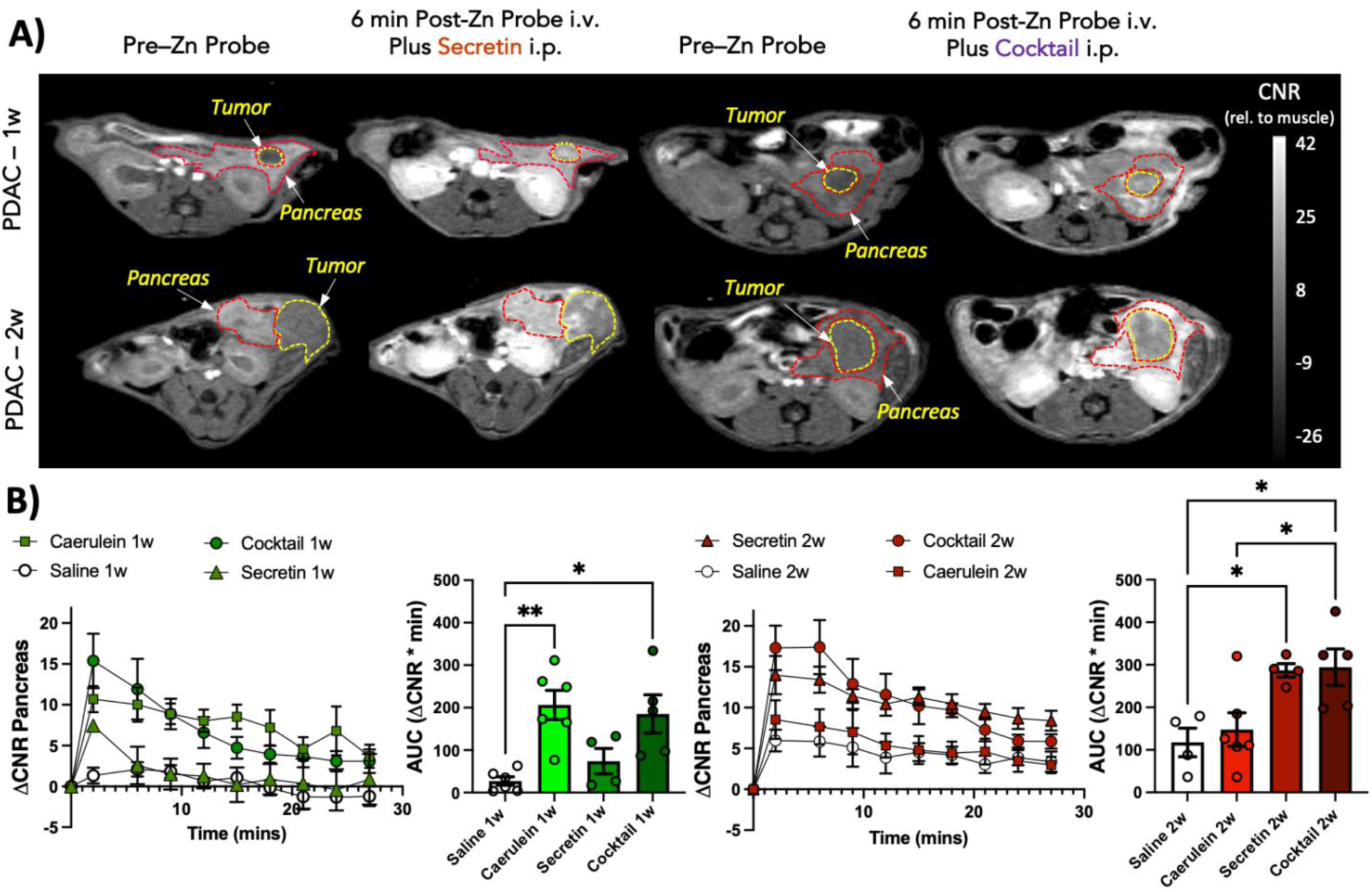
A cocktail of secretin and caerulein secretagogues maximizes zinc secretion in PDAC. **A)** *In vivo* T1-weighted MR images before and 6 minutes after 0.07 mmol/kg i.v. GdL injection and either 50 µl Secretin or 10 µg/kg secretin plus caerulein cocktail i.p. injection in 1-week and 2-week PDAC mice. Regions of interest were drawn to outline the tumor (yellow) and the surrounding pancreas (red). **B)** Dynamic change in CNR (△CNR) of tumor-bearing pancreas or tumor ROIs relative to muscle signal over 27min. Integrated change in CNR over the imaging period (Area under the curve, AUC) shows maximal enhancement in the pancreas after stimulation with secretin plus caerulein administered as a cocktail. Error bars are SEM. Statistical significance was determined by one-way ANOVA with Tukey test for multiple comparisons. *p<0.05, ** p < 0.01, *** p <0.001, **** p<0.0001.

We evaluated the expression of zinc transporters in the tumor-bearing pancreas of animals with 1-week and 2-week-old tumors. Immunohistochemistry was performed to stain against ZIP3, ZIP4, and ZNT1. We segmented the pancreas and the tumor as seen in **Figure 4A** and selected ROIs for each tissue, representative pancreas ROIs are shown in **Figure 4B**. In the pancreas of 1-week PDAC animals there is significantly higher ZIP3 expression compared to the pancreas of healthy controls, while the pancreas of 2-week PDAC animals shows significantly higher import and export transporter expression than the healthy pancreas (ZIP3: 16 ± 5 % vs. 2 ± 1 %, p<0.0001; ZIP4: 15 ± 9 % vs. 2 ± 1 %, p<0.0001; 6 ± 4 vs. 2 ± 1, p < 0.001). There is a significantly higher expression of zinc transporters (ZIP3, ZIP4, and ZnT1) in the pancreas of week-2 PDAC animals compared to week-1 animals (ZIP3: 16 ± 6 vs. 5 ± 3, p < 0.0001; ZIP4: 15 ± 9 vs. 3 ± 2, p< 0.001; ZnT1: 5 ± 2 vs. 8 ± 6, p<0.05) **Figure. 4C**).

**Figure 4.**
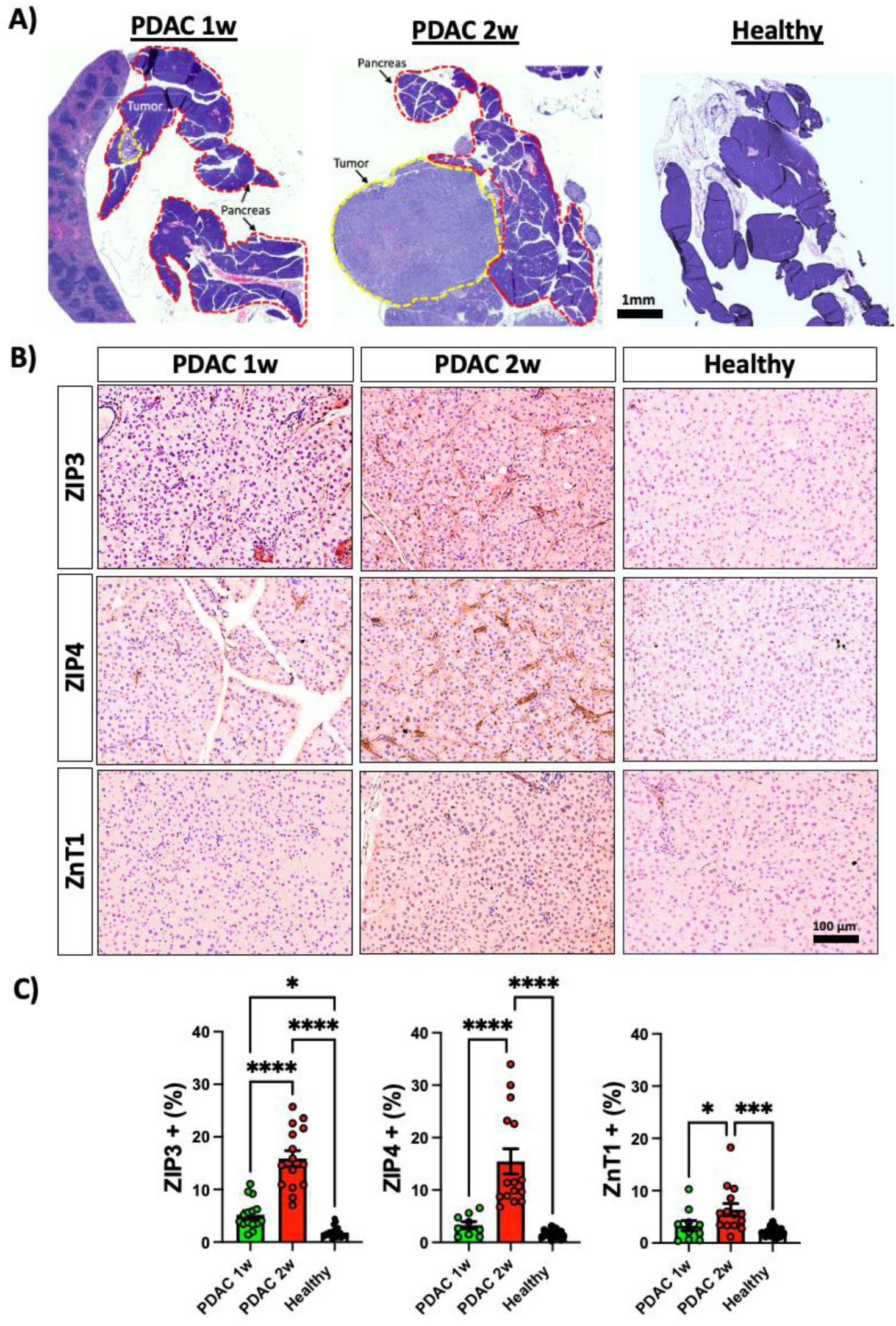
Zinc transporter expression in the pancreas of healthy mice and mice bearing PDAC tumors for 1- or 2- weeks. **A)** H&E staining of PDAC mouse pancreas bearing a tumor for 1 week and 2 weeks. The surrounding pancreas tissue was delineated and analyzed. **B)** Immunohistochemistry of pancreas tissue surrounding 1-week and 2-week PDAC tumors. **C)** ROIs were drawn in the pancreas to quantify zinc transporter expression. The level of expression was calculated as positive transporter staining cells relative to total cells in each ROI drawn in 1-week, 2-week PDAC, and healthy pancreas. Error bars are SEM. Statistical significance was obtained by Two-way Anova corrected by a Tukey test for multiple comparisons. ** p < 0.01, *** p <0.001, **** p<0.0001.

### The exocrine pancreas does not exhibit stimulated zinc hypersecretion in a surgical model of pancreatitis

To determine if our imaging technique may differentiate PDAC from non-cancer related pancreatitis we stimulated zinc secretion in a surgical model of acute pancreatic inflammation. Surgical ligation of the pancreatic duct results in an obstruction of pancreatic juice drainage out of the tail region of the pancreas, therefore initiating tissue damage with progressive acinar cell atrophy, tissue necrosis, metaplastic duct formation, and immune cell infiltration. (37) In this model, and for this experiment, the pancreatic duct is surgically ligated in the neck area of the pancreas. SSZS-MRI was performed 3 days and 7 days post ligation. **(Figure 5A *top*).** Hematoxylin and Eosin (H&E) staining show that seven days post ligation surgery, the tail (ligated part of the pancreas) is injured, inflamed, and small metaplastic ducts are formed. In contrast, the pancreas head (un-ligated part of the pancreas) remains intact and healthy **(Figure 5A *bottom*)**. We analyzed the expression of ZIP3, ZIP4, and ZNT1 in the ligated tail, un-ligated head, and sham pancreas tissue (**Figure 5B**). Zinc importers, ZIP3, and ZIP4 expression increased significantly in the ligated tail of the pancreas compared to the head of the pancreas and sham pancreas, as shown by IHC. Zinc export transporter expression, ZNT1, also increased significantly in the ligated pancreas compared to the un-ligated healthy tissue **(Figure 5C)**. However, it is important to note that the level of expression is low (∼2%) in inflamed pancreas compared to pancreas containing PDAC (∼15%).

**Figure 5.**
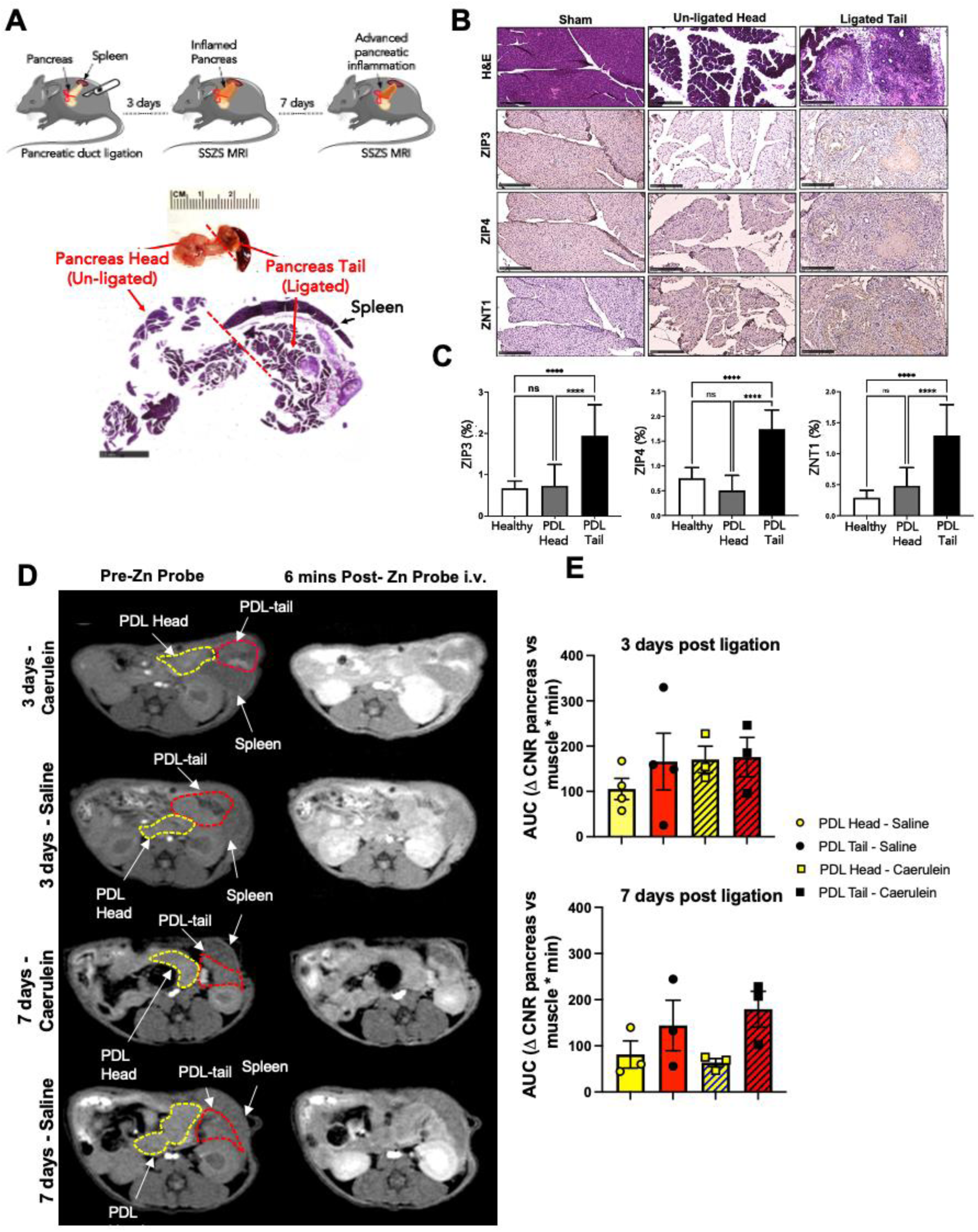
SSZS MRI of mice with ligated pancreas duct as a model of pancreatitis. **A)** Surgical procedure and imaging schedule. **B)** H&E and IHC staining against ZIP3, ZIP4, and ZNT1 of sham animals, un-ligated head portion of the pancreas, and ligated tail of the pancreas showing severe inflammation**. C)** Quantification of zinc transporter expression in tissue sections. **D**) *In vivo* SSZS - MRI of animals after 3 days and 7 days post pancreatic duct ligation. The ligated portion of the pancreas is outlined in red and labeled as PDL-Tail and the un-ligated portion is outlined in yellow and denoted as PDL-Head. Animals either received saline or caerulein at each timepoint. **E)** Calculated area under the curve (AUC) of each pancreas portion (tail, and head) where the signal change relative to muscle over 27 minutes was integrated and compared for each caerulein or saline group. (N = 3 each). Error bars are SEM. Statistical significance was determined by one-way ANOVA with Tukey test for multiple comparisons. *p<0.05, ** p < 0.01, *** p <0.001, **** p<0.0001.

The ligated pancreas was identified by MRI first by examining the anatomy in both axial and coronal T_2_-weighted MRI scans. The ligated tail (PDL-tail) was identified as the ligated portion closest to the spleen, and the healthy head (PDL-head) was located as the region of the pancreas directly adjacent to the ligation (**Figure 5A *bottom* and 5D**). Once the regions were identified, axial T_1_-weighted scans were obtained to encompass both ligated portions. **Figure 5D** shows the pre-injection scans where the PDL-tail and PDL-head were drawn as red and yellow ROIs, respectively. The T_1_-weighted scans 6 minutes post administration of Zn probe and either saline or caerulein indicate that there is a slight increase in SSZS in the ligated pancreas versus the adjacent differences amongst groups at either three days or seven days post pancreatic duct ligation. Additionally, we used a previously reported molecular probe that “turns on” in the presence of reactive oxygen species (ROS) found in tissue with high inflammatory cell infiltrate, FePyC3A (38). After administering 0.3 mmol/Kg, we observed a significant increase in MRI signal in the ligated tail compared to the “healthy” head (**Supplemental Figure 3A *left***). This enhancement was quantified by integrating the ΔCNR over time for each one of these tissues. **Supplemental Figure 2 A *right*** shows that there is a 2.5-fold increase in integrated ΔCNR in the PDL tail compared to the PDL head consistent with increased ROS due to the inflammatory state of the PDL tail. The pancreas was then harvested and both ligated tail and unligated head (healthy) were stained against myeloperoxidase (MPO). The portion of cells stained positively for MPO was quantified and represented as MPO (%), **Supplemental Figure 2B *left*** shows immunohistochemistry staining illustrating significant levels of inflammatory cell infiltrate in the ligated tissue, and **Supplemental Figure 2B *right*** illustrates the increased MPO (%) staining in the ligated tail compared to the un-ligated head of the pancreas.

Combining these two different molecular imaging techniques we were able to show that SSZS-MRI only promotes signal change in the tumor-bearing pancreas and not in pathologically benign pancreatitis.

### SSZS-MRI detects response to KRAS G12D inhibitor treatment

MRTX1133, a small-molecule inhibitor targeting the KRASG12D mutation, has recently shown significant potential in treating PDAC (39). The orthotopic PDAC model used here involves the implantation of cells that carry the G12D mutation, and so we tested the ability to detect response to KRASG12D inhibitor treatment by SSZS-MRI. **Figure 6A** shows the treatment schedule, imaging, and tissue collection plans for these experiments. We treated PDAC animals with either MRTX1133 or vehicle (Captisol) i.p. BID for 17 days following previously reported protocols (39) and performed SSZS-MRI 3-5, 10-12, and 16-17 days post treatment initiation. Response to treatment was assessed by measuring the tumor volume in anatomical T_2_-weighted MRI scans obtained at every imaging time point. We found that the mean tumor volume for animals receiving vehicle was significantly higher than MRTX1133 groups even at the first days of treatment initiation (vehicle: 247 ± 68 mm^3^ vs. 3-5d: 42 ± 22 mm^3^) (**Figure 6B**). In one week, there was a 4-fold increase in tumor size, which prompted us to sacrifice the vehicle group animals earlier and collect the tissue after 10-12 days for ex vivo analyses. Conversely, in the MRTX1133 group, tumor growth was halted at the first days of treatment initiation and in most cases continued to decrease in size shrinking the tumor for the duration of the treatment schedule (16-17 days). It is noteworthy that upon close inspection of the tumor volumes between cocktail-stimulated and saline groups a trend emerged in which the cocktail-stimulated group appeared to have smaller tumor volumes compared to tumors in the saline-stimulated groups (**Figure 6B**).

**Figure 6.**
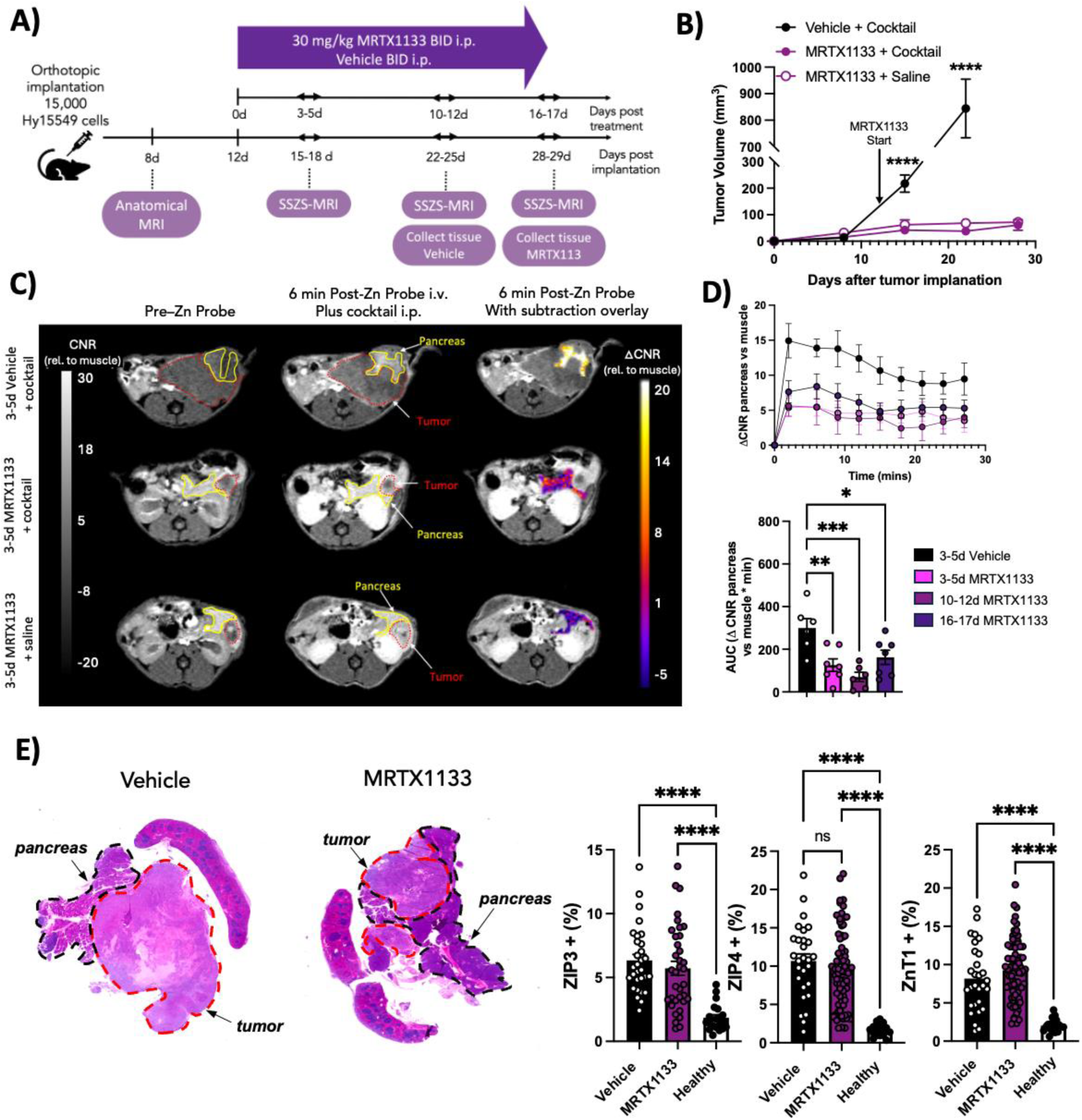
Stimulated zinc secretion MRI detects response to KRASG12D inhibitor, MRTX1133, treatment in PDAC. **A)** Experimental plan including tumor cell implantation, treatment regimen, imaging and tissue collection time points. **B)** Mean tumor volume measured from T_2_-weighted MRI anatomical scans over time. **C)** T_1_-weighted gradient echo images of animals 3-5 days post treatment with *(top two rows)* MRTX1133 and Vehicle prior to and 6 min after receiving 0.07 mmol/kg Zn Probe iv. and 10 µg/kg secretagogue cocktail i.p. or *(bottom row)* saline i.p. as control. **D)** *(top)* Dynamic ΔCNR of tumor bearing pancreas vs. muscle. *(bottom)* Area under the ΔCNR vs. time curve for animals receiving vehicle control (N = 6) or MRTX1133 treatment for 3-5, 10-12, or 16-17 days (N = 7). **E)** H&E of representative Vehicle, and MRTX1133 treated tissue collected 10-12d and 16-17d post treatment, respectively. Quantified zinc transporter (ZIP3, ZIP4, ZnT1) expression was obtained from pancreas tissue for Vehicle, MRTX1133 treated, and Healthy animals. Each value is an ROI within the segmented pancreas (black outline). Error bars are SEM. Statistical significance was determined by one-way ANOVA with Tukey test for multiple comparisons. *p<0.05, ** p < 0.01, *** p <0.001, **** p<0.0001.

To assess the ability to detect treatment response based on stimulated zinc secretion we performed SSZS-MRI. **Figure 6C** shows T1-weighted MRI scans of animals treated with MRTX1133 vs. Vehicle before and 6 min after zinc probe and secretagogue stimulation with cocktail (secretin + caerulein) or with saline. In the cocktail-stimulated vehicle-treated group, the tumor size is dramatically larger, and the surrounding pancreas is significantly more enhanced compared to the cocktail-stimulated and non-stimulated MRTX1133 treated groups. Serial MRI scans acquired for 27 min after injections show that there is significant increase in pancreas ΔCNR in the vehicle group compared to MRTX1133 groups (**Figure 6D**). A 3-fold reduction in AUC (**Figure 6D *bottom panel***) was observed between vehicle and treated groups (3-5d vehicle: 149 ± 17 ΔCNR *min vs. 3-5d: 120 ± 23 ΔCNR *min, 10-12d: 102 ± 33 ΔCNR *min and 16-17d: 162 ± 24 ΔCNR *min, p<0.0001). Saline-stimulated groups did not exhibit differences in MRI signal in the pancreas during MRTX1133 treatment (**Supplemental Figure 3A**).

To study the zinc transporter expression in this treatment-response model, we collected the tissues on days 10-12 for vehicle group, days 16-17 post treatment for MRTX1133 group and compared them to healthy pancreas. The pancreas was segmented and separated from the tumor as shown in **Figure 6E left**. The import and export transporters ZIP3, ZIP4, and ZnT1 were quantified from 5 ROIs drawn throughout the segmented pancreas tissue. All transporters remained significantly upregulated compared to healthy animals however, there was no difference in expression between vehicle and MRTX1133 treated groups as shown on **Figure 6E right**.

### SSZS-MRI detects PDAC recurrence after MRTX1133 treatment withdrawal

We then tested our imaging technique in a model of PDAC recurrence after MRTX1133 treatment withdrawal. As seen in **Figure 7A**, we orthotopically implanted PDAC cells in the mouse pancreas and assigned them to two groups: **1)** secretagogue cocktail stimulation and **2)** saline stimulation. A pre-treatment anatomical MRI was performed to assess successful tumor implantation. Treatment with 30 mg/kg MRTX1133 BID started on day 9 until day 21, SSZS MRI was performed immediately before treatment withdrawal, and after 1-3 days, and 11-12 days post withdrawal. Tissues were collected on day 18-19 post-treatment withdrawal. Tumor burden was measured by outlining the tumor volume from T_2_-weighted MRI scans and as expected, after treatment withdrawal, the tumor recurs and grows gradually as seen in **Figure 7B**. Interestingly, during treatment withdrawal we observed that the animals stimulated by secretagogue cocktail exhibited slower cancer recurrence (smaller tumor volumes) compared to saline-stimulated animals (larger tumor volume). This divergence in growth rates was detected as early as 1-3 days post treatment withdrawal (secretagogue cocktail group: 83 ± 28 mm^3^ vs. saline group: 131 ± 40 mm^3^). **Figure 7C *(top row)*** illustrates pre-MRTX1133 withdrawal scans where the change in pancreas signal was low consistent with response to MRTX1133 treatment. During cancer recurrence, we see that the more the tumor grows after treatment withdrawal the higher the SSZS MRI enhancement observed in the pancreas. **Figure 7D** shows the ΔCNR in the pancreas over the entire imaging period, by calculating the AUC from the dynamic scans we found that there is a statistically significant difference between the AUC in the pre-withdrawal scans, 1-3, and 11-12 days post-MRTX1133 withdrawal (Pre-WD: 68 ± 43 vs. 1-3d WD: 128 ±64, and vs. 11-12d WD: 215 ±122 ΔCNR*min, p< 0.05) (**Figure 7C**). The saline stimulated animals exhibited no statistical difference between groups (**Supplemental Figure 3B**). At the end of the study, we collected the tissue to perform histology and immunohistochemistry to evaluate the expression of zinc transporters (ZIP3, ZIP4, and ZNT1). **Figure 7E** illustrates the difference in tumor size at the end of MRTX1133 treatment (pre-withdrawal) and 18-19 days with no treatment at all. The tumor was delineated, and the surrounding pancreas was segmented for quantification of zinc transporter expression. Only ZIP3 exhibited significant increase after MRTX1133 treatment withdrawal (MRTX1133 Pre-WD: 6 ± 3 % vs. MRTX1133 WD: 8 ± 3%) (**Figure 7E**).

**Figure 7.**
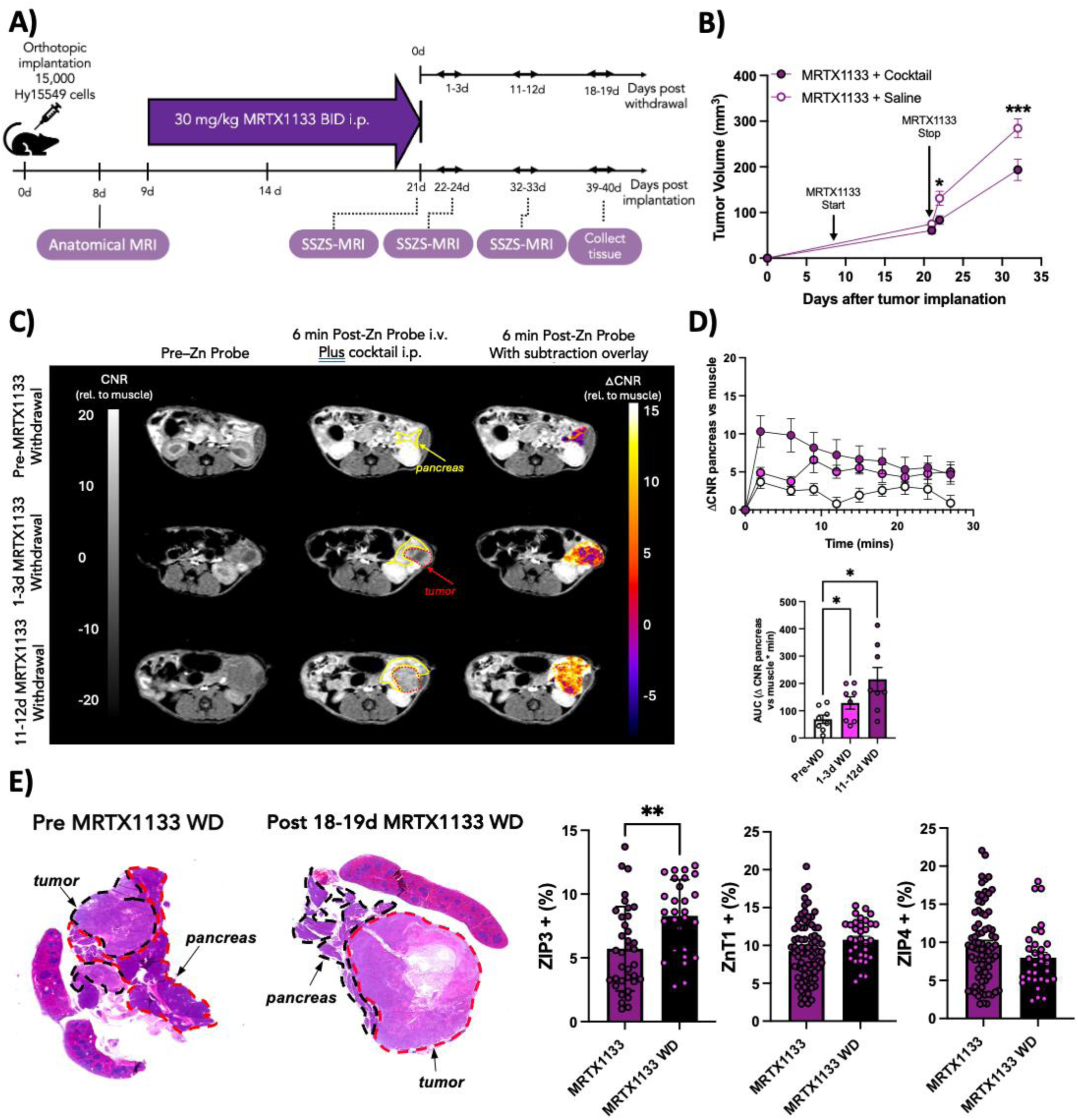
Stimulated zinc secretion MRI detects cancer recurrence after KRASG12D inhibitor treatment withdrawal. **A)** Experimental plan including tumor cell implantation, treatment regimen and withdrawal, imaging and tissue collection time points after withdrawal. **B)** Mean tumor volume of PDAC animals at the end of MRTX1133 treatment, 1-3d, and 11-12d after treatment withdrawal. Animals were scanned by SSZS-MRI and received either saline or cocktail as a secretagogue. **C)** T_1_-weighted gradient echo MR images of pre-MRTX1133 treatment withdrawal, 1-3 days post-treatment withdrawal, and 11–12-day post-treatment withdrawal before and 6 min after 0.07 mmol/kg zinc probe iv. injection while stimulated with cocktail i.p. (N=8) **D)** *(top)* Dynamic ΔCNR of tumor-bearing pancreas tissue vs. muscle. *(bottom)* The area under the curve of ΔCNR over the entire imaging period. **E)** H&E of representative pre-withdrawal, and 18-19d post withdrawal tissue. Quantified zinc transporter (ZIP3, ZIP4, ZnT1) expression was obtained from pancreas tissue for Pre-withdrawal and 18-19d post treatment withdrawal. Each value is an ROI within the segmented pancreas (black outline). Error bars are SEM. Statistical significance was determined by one-way ANOVA with Tukey test for multiple comparisons. *p<0.05, ** p < 0.01, *** p <0.001, **** p<0.0001.

## DISCUSSION

PDAC remains an unsurmountable challenge in oncology due to its poor prognosis and the difficulty of early detection. Traditional imaging techniques and biomarkers often fail to provide timely and precise information necessary for optimal clinical management. This study addresses the critical need for improved diagnostic and monitoring tools for PDAC, focusing on the potential of SSZS-MRI to meet these challenges.

Our study demonstrated that the tumor bearing pancreas exhibits secretagogue-stimulated zinc hypersecretion compared to the healthy and pathologically benign pancreas as shown in healthy, ligated pancreas, and PDAC mice of two different genetic backgrounds (C57Bl6 and FVB). The ability of SSZS-MRI to detect these changes underscores its potential for more accurate diagnosis. Furthermore, we found that zinc secretion can be maximized by using multiple secretagogues (caerulein and secretin) administered as a cocktail. Caerulein, a cholecystokinin analog, preferentially stimulates secretion of acinar cells while secretin promotes secretion from ductal cell types in the pancreas (40) and as such administering them as a cocktail results in maximal stimulated secretion. Importantly, SSZS-MRI with secretagogue cocktail detected responses to KRASG12D treatment as early as 3-5 days post-treatment initiation and identified PDAC recurrence as early as 1 day after KRASG12D treatment withdrawal. These findings highlight the timeliness of SSZS-MRI in monitoring treatment efficacy, enabling early intervention and potentially reducing unnecessary cytotoxic exposure. A distinctive advantage of pancreas SSZS-MRI is its ability to gather information regarding cancer occurrence from the pancreas itself, rather than relying solely on tumor imaging. This is particularly important as pancreatic tumors can be elusive and difficult to locate due to their small size.(41) Our study found that dysregulation of zinc transporters in human and mouse PDAC tissues as a hallmark of the disease. This zinc transporter dysregulation offers a functional imaging approach through pancreatic zinc secretion patterns that if found dysfunctional it could prompt further careful examination to identify occult small lesions.

The use of secretagogues in conjunction with SSZS-MRI not only enhances imaging capabilities but also appears to boost treatment response and hinder recurrence. Our study found that animals receiving secretagogue stimulation exhibited improved treatment response and slower cancer recurrence, as measured by tumor volume during KRASG12D inhibitor treatment and after withdrawal. Given the multifactorial effects that KRASG12D inhibitors have at reprograming the tumor micro- and macroenvironment (42,43) further studies are necessary to elucidate the mechanisms of improved treatment response and delayed cancer recurrence. Nevertheless, this dual function of secretagogues—both as zinc imaging enhancers and potential therapeutic adjuvants—opens new avenues for integrated diagnostic and therapeutic approaches. The controlled stimulation of pancreatic secretion could improve detection and treatment efficacy by sensitizing tumors to subsequent therapeutic interventions.

Other imaging approaches that aim at detecting PDAC occurrence early include the use of fibroblast activation protein (FAP)-targeted PET probes, which have shown promise in detecting PDAC,(44) however its applicability to detect treatment response has not yet been fully elucidated. Alternatively, an emerging field of study in metabolic imaging uses hyperpolarized ^13^C pyruvate to image the metabolic conversion of pyruvate to lactate in solid PDAC tumors. This technology has been tested in small cohort of PDAC patients (N=6) showing promising results where it was observed that there was altered pyruvate metabolism with increased lactate or reduced alanine in primary tumors.(45) However, this technology is limited by the rate of loss of polarization and requires completely new infrastructure and imaging workflows that are costly raising cost-benefit concerns. Therefore, an imaging technique that could be easily incorporated into the current imaging workflow and combine the biological functional information of the pancreas with the non-invasive and anatomical qualities of MRI/MRCP would be extremely valuable.

The clinical implications of SSZS-MRI are profound, with potential applications extending beyond PDAC to other malignancies characterized by zinc dysregulation, such as prostate and breast cancer (13,27,46,47). The non-invasive nature of SSZS-MRI, coupled with its ability to provide both anatomical and functional information, makes it an attractive option for routine clinical use. Additionally, the technology could be integrated into existing imaging workflows, facilitating widespread adoption without significant infrastructural changes. Early detection capabilities of SSZS-MRI could lead to improved survival rates by enabling timely surgical interventions and more personalized treatment regimens.

Despite the promising results, there are limitations to this study. 1) The experimental set up of this study requires that in order to assess zinc flux two separate groups of mice were required to receive secretagogue and saline, respectively due to concerns regarding injection volume overload. In humans this would be simplified as both saline and secretagogue could be administered sequentially. 2) In this study we only tested our SSZS-MRI technology on male mice given the slight increased incidence of PDAC occurrence in males versus females, however the effect of sex as a biological variable needs to be evaluated. 3) In addition, further validation of other zinc transporters is needed as there may be additional candidates of the ZIP and ZnT families that may show upregulation and malignant specificity during cancer onset and progression and importantly that may be more sensitive to treatment response with KRASG12D inhibitors.

In conclusion, SSZS-MRI represents a significant advancement in pancreatic cancer imaging, offering enhanced sensitivity and specificity for early detection and treatment monitoring compared to non-functional contrast-enhanced MRI. By leveraging zinc homeostasis dysregulation in the organ as a whole, this technology provides valuable molecular insights that complement traditional imaging modalities. The potential adjuvant therapeutic applications of secretagogues further enhance the clinical utility of SSZS-MRI. While challenges remain, the potential clinical benefits of SSZS-MRI warrant further investigation and development, aiming to improve outcomes for patients with PDAC and other zinc-dysregulated cancers.

## METHODS

### GdL zinc probe and secretagogue preparation

Gd-based zinc probe, GdL, was obtained from VitalQuan, LLC (Dallas, TX) in multiple batches. 50 mg each dissolved in 0.5 mL water. The compound was formulated for each animal experiment to a 30 mM stock in saline. The dose was 0.07 mmol/kg and was administered from stock as 2 µL/g_mouse_.

Secretin and caerulein were obtained from Fisher Scientific and formulated in saline to administer a dose of 10 µL/kg of each or as a cocktail.

### Human samples

All human samples were acquired with consent and in accordance with the protocols approved by the Massachusetts General Hospital Institutional Review Boards (IRBs).

### Immunohistochemistry (IHC)

The pancreas and tumor tissue were fixed with 10% formalin for 48 hours prior to paraffin embedding. Sections were cut at 7 µm thickness. Representative sections were stained with Hematoxylin & Eosin (H&E). Antibodies targeting ZIP3 (1:100, Abcam, Cambridge, MA, USA), ZIP4 (1:100, Abcam, Cambridge, MA, USA), ZNT1(1:100, Abcam, Cambridge, MA, USA) were used to perform immunohistochemistry according to our previously published protocols. (34) Images were captured with NanoZoomer-SQ Digital slide scanner (Hamamatsu Photonics, Japan). All image analyses were obtained using ImageJ software and performed by a blinded reviewer (National Institutes of Health (NIH)).

### Mouse model and PDAC implantation

All animals received humane care based on the NIH Guide for the Care and Use of Laboratory Animals and procedures were approved by the Massachusetts General Hospital Institutional Animal Care and Use Committee (IACUC). Experiments were designed in accordance with the Animal Research: Reporting in vivo experiments (ARRIVE) guidelines. 8-10 weeks old male C57BL/6 and FVB mice were purchased from Charles River Laboratories (Wilmington, MA). Mice were fed regular chow and housed in a light and temperature-controlled animal facility. Animals were randomly assigned to experimental groups. Tumor implantation has been performed according to our previously published protocols. (34,48) In brief, Hy15549 (*Kras^G12D^* and *p53^+/−^*) and Han4.13 (*Kras^G12D^*and *p53^+/−^*) murine PDAC cell lines were isolated from primary pancreatic tumors generated from Ptf1-Cre; LSL-KRAS^G12D^; p53^Lox/+^ mice. Hy15549 cells are syngeneic with the C57BL/6 mouse strain, and Han4.13 cells are syngeneic with the FVB mouse strain. Cells were cultured in Dulbecco’s Modified Eagle’s Medium (DMEM) (high glucose 4.5 g/L) (Gibco, Grand Island, NY, USA) supplemented with heat-inactivated 10% FBS (Sigma-Aldrich, St. Louis, MO, USA) and 1% Penicillin /Streptomycin. Cells were detached with 0.25% trypsin for 2-3 minutes at 37°C. The suspended cells were centrifuged at 1300 RPM x 5 min, resuspended in 1 mL of DMEM media, and counted using the Countess Automated Cell Counter (Life Technologies, Carlsbad, CA, USA). Both Hy15549 and Han4.3 cells were resuspended at a concentration of 10^3^ cells/*μ*L in Matrigel and kept on ice until the time of injection. 10^4^ cells/10 µL were injected orthotopically in the pancreas. Before the cell injection, fur was shaved in the incision region using an electric clipper. The surgical field was sterilized with betadine. A 0.5 cm left flank incision was made, and the pancreas and spleen were extracorporealized. An Insulin syringe was used to inject cells into the body of the pancreas. Within 1 minute after cell injection, Matrigel solidifies. The pancreas and spleen were placed carefully back in the body without imposing pressure on the injection site to avoid leakage. The muscle was closed with 5-0 Polypropylene sutures, and the skin was closed with surgical staples. Mice were given Buprenorphine (0.05 – 0.1 mg/kg) analgesia by subcutaneous flank injection every 6 hours for pain control. Mice were monitored daily for any distress and pain for up to 2 weeks. The tumor and pancreas were extracted after each MRI for histological evaluation.

### Pancreatic duct ligation as a model for pancreatitis

Pancreatic duct ligation was performed according to published protocols. (37,49) 8 weeks male mice were anesthetized with an intraperitoneal injection of ketamine (100 mg/kg) and xylazine (10 mg/kg). Fur was shaved in the lower back of the animal’s left side. The surgical field was sterilized with betadine and was allowed to dry for 30 seconds. A 1 cm incision was made. The spleen and pancreas were gently taken out and exposed. The neck area was ligated using a 6-0 Prolene suture. Next, the pancreas and spleen were gently put back, and the wound was closed in two layers: muscle with running 6-0 Prolene suture, skin with surgical clips. Mice were given a single bolus of 500 µL of PBS by subcutaneous flank injection for management of the ensuing acute pancreatitis. Mice were given Buprenorphine (0.05 – 0.1 mg/kg) analgesia by subcutaneous flank injection every 6 hours for pain control. Mice were monitored daily for morbidity, evidenced by decreased grooming, hunched posture, and reduced movement.

### *In vivo* MRI and image analysis

Animals were anesthetized with 1-2% isoflurane with body temperature maintained at 37 °C. The tail vein was cannulated for intravenous delivery of zinc probe GdL. Caerulein, cocktail (secretin+ caerulein), or saline were administered intraperitoneally while the animal was positioned in the scanner in a custom-designed cradle. Imaging was performed at 4.7 T using a small-bore animal scanner (Bruker, Billerica, MA). 2D axial and coronal T_2_-weighted Turbo Rapid Imaging with Refocused Echoes (Turbo RARE) scans were obtained to locate the tumor and pancreas for dynamic T_1_-weighted MRI. The turbo RARE parameters were as follows: TE/TR= 27/1360.21 ms, echo train length = 8, flip angle = 90, FOV=33x33 mm, matrix=192x192, slice thickness=1mm, Averages = 4. Once the slice package that optimally covers the tumor and surrounding pancreas was created, baseline axial and coronal 2D Fast Low Angle Shot (FLASH) images were acquired before and continuously for 30 minutes after injection of 10 µg/kg caerulein intraperitoneally, and 0.07 mmol/kg GdL intravenously 5 minutes after. 2D FLASH image acquisition parameters were TE/TR = 2.93/152 ms, flip angle FA=60°, FOV=33x33 mm, matrix=140x140, and slice thickness = 1.0 mm. The animals were sacrificed upon completion of scans, and tissue was prepared for histology.

Data analysis included the calculation of the contrast-to-noise ratio of the pancreas region of interest (ROI) relative to back muscle ROI, and an ROI placed outside the body encompassing air (i.e., 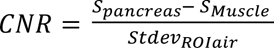). The differential (postinjection – preinjection) was further calculated and represented as change in CNR (ΔCNR). The area under the ΔCNR vs. time curve (AUC) was calculated for each treatment group using GraphPad Prism Software (GraphPad Software, La Jolla, CA, USA). Tumor burden was evaluated by outlining tumor tissue on each slice. The area of that ROI was measured and multiplied by the slice thickness (1 mm). Overall tumor volume (expressed in mm^3^) was computed by adding all the volume calculations for each tumor-containing slice.

### KRASG12D inhibitor, MRTX1133, treatment

MRTX1133 was provided by Mirati Therapeutics and prepared using 10% research-grade Captisol (CyDex Pharmaceuticals) in 50 mmol/L citrate buffer pH 5.0 (Teknova, Q2443). The compound was made weekly and was stored at 4°C while protected from light. After tumor implantation, mice were monitored daily for signs of tumor growth. Tumors were allowed to develop following the growth and treatment schedule outlined in **Figure 6A**, on day 11, the animals began receiving twice-daily intraperitoneal (IP) injections of MRTX1133 at a dose of 30mg/kg for the treatment groups, while the Vehicle group received Captisol (30mg/kg), continuing until day 28. For the cancer recurrence model, following the schedule outlined in **Figure 7A**, treatment started on day 9 and continued until day 21, after which mice were withdrawn from the treatment.

### Statistical analysis

Results are expressed as the mean ± 1 SEM unless otherwise noted. One-way ANOVA followed by post-hoc Tukey tests with 2-tailed distribution were performed to analyze data among groups of 3 or more. To evaluate the effects of a variable according to the levels of two other variables we used a Two-way ANOVA followed by post-hoc Tukey test. The student’s t-test compared data between two experimental groups. All tests were performed 2-sided, and a significance level of p < 0.05 was considered statistically significant. All statistical analyses were performed using GraphPad Prism software (GraphPad Software, La Jolla, CA, USA).

