## Supplementary Figures 1 - 3 for "Molecular Magnetic Resonance Imaging of Dysregulated Zinc Secretion Detects Early Pancreatic Ductal Adenocarcinoma Lesions and Response to KRASG12D Inhibitor Treatment"

**SUPPLEMENTARY FILES**

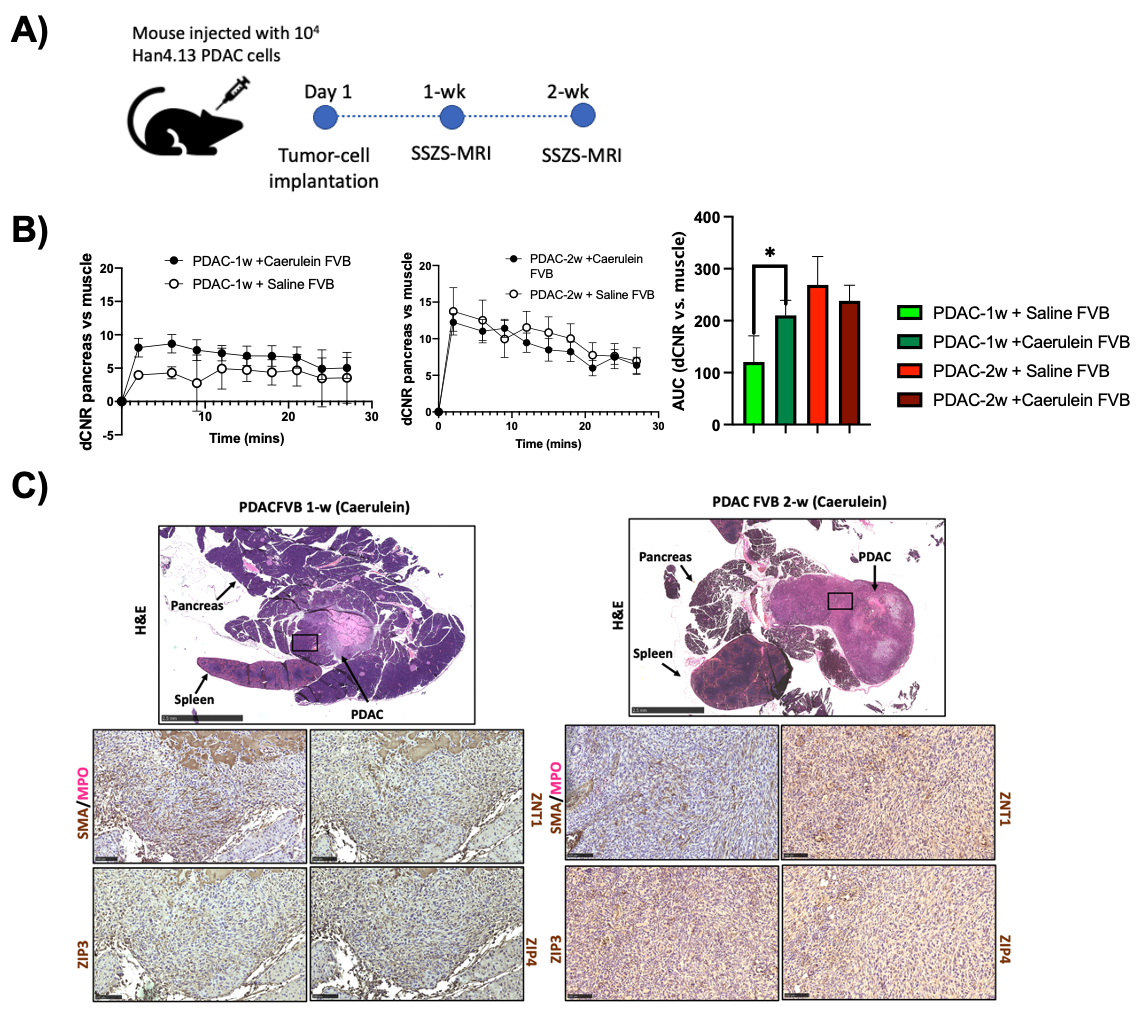

**Figure 1S. Stimulated zinc secretion in PDAC mouse of FVB genetic background. A)** Orthotopic PDAC cell implantation and imaging schedule**. B)** (left) Change in contrast to noise ratio (ΔCNR) after zinc probe and caerulein or saline for PDAC animals bearing Han4.13 cell tumors for 1 week and 2 weeks. (right) The area under the curve of pancreas ΔCNR over the entire 27-minute imaging period for all groups. **C)** Tissue H&E showing the presence of 1-week old and 2-week-old tumors (Arrow pointing PDAC) and IHC panels showing immunostaining against ZIP3, ZIP4, ZNT1, and SMA/MPO as inflammatory markers. Error bars are SEM. Statistical significance was determined by one-way ANOVA with Tukey test for multiple comparisons. *p<0.05.

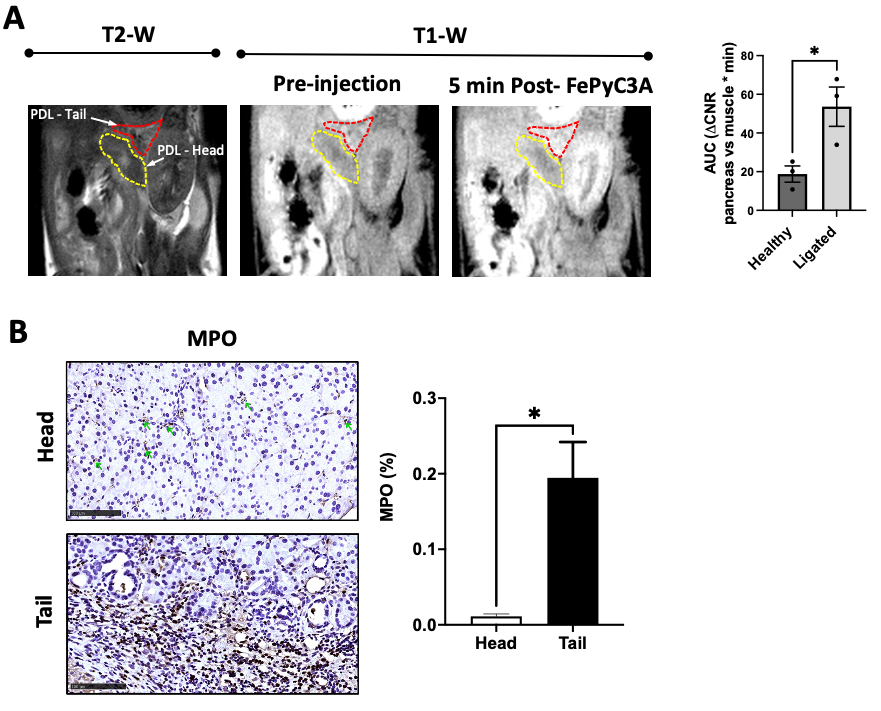

**Figure 2S. Inflammation in the pancreas with pancreatic duct ligation. A)** *(Left)* *In vivo* MRI of the inflamed pancreas. The ligated tail and un-ligated head of the pancreas were identified by anatomical T_2_-weighted MRI scans. Molecular imaging of inflammation in the pancreas was obtained by administering the ROS-active probe FePyC3A. The tail of the pancreas exhibited differential enhancement compared to the head of the pancreas indicating the presence of ROS as a hallmark of a highly inflammatory state in the ligated pancreas. *(Right)* Calculated area under the curve (AUC) of head and tail of the pancreas in mice after receiving the inflammation sensitive probe FePyC3A. **B)** (*Left*) IHC staining against myeloperoxidase (MPO) for both ligated pancreas portions, head, and tail. (*Right*) Blinded quantification of MPO expression in tissue sections for both pancreatic tissue portions, head, and tail. Error bars are SEM, Statistical significance was obtained by two-tailed t-test. * P < 0.05

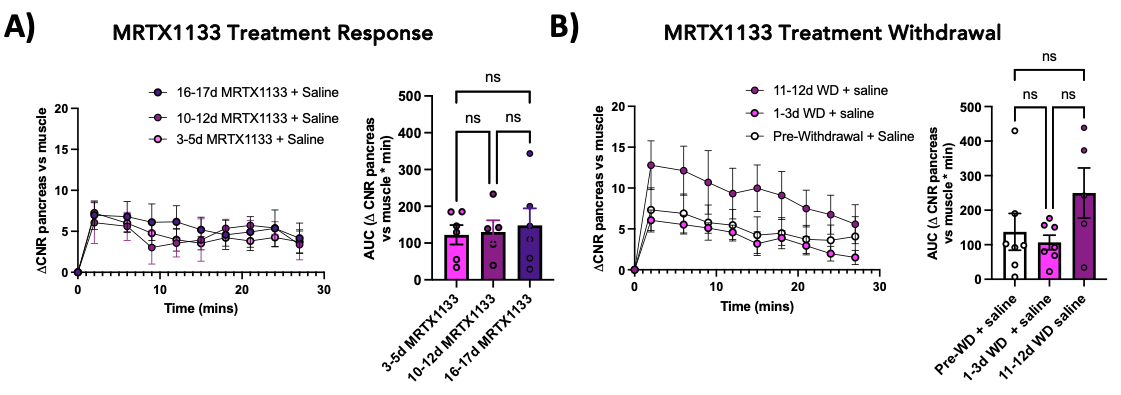

**Figure 3S. MRI of unstimulated animals on MRTX1133 treatment and after MRTX1133 treatment withdrawal.** *(Left)* Quantified dynamic change in CNR of tumor-bearing pancreas tissue vs. muscle over time. (*Right*) Area under the change in CNR over the entire imaging period while on **A)** MRTX1133 treatment and receiving saline as secretagogue control or after **B)** MRTX1133 treatment withdrawal. Error bars are SEM. Statistical significance was determined by one-way ANOVA with Tukey test for multiple comparisons.
